# Default Mode and Dorsal Attention Network functional connectivity associated with alpha and beta peak frequency in individuals

**DOI:** 10.1101/2023.02.19.529136

**Authors:** Vaibhav Tripathi, David C Somers

## Abstract

Alpha- and Beta-frequency oscillatory waves are evident in resting-state electro- and magneto-encephalography (EEG, MEG). Higher peak frequencies for Alpha and Beta are associated with greater cognitive health. Aging slows down the alpha and beta waves which are also affected by mental states, disorders such as ADHD, and sleep deprivation. Functional magnetic resonance imaging (fMRI) of resting-state brain activity reveals anti-correlation of very low frequency (< 0.1 Hz) fluctuations between two distributed cerebral cortical networks, the Dorsal Attention Network (DAN) and the Default Mode Network (DMN). DAN activation is related to attentional demands in extrinsic tasks, whereas DMN is associated with mind wandering, episodic memory retrieval, and intrinsic processing. Prior research has found that higher DAN-DMN anticorrelation is related to greater cognitive and mental health. Here, we investigated whether these two measures of cognitive and mental health are related to each other within individuals. We investigated resting-state Functional Connectivity (rsFC) between the DMN and DAN using fMRI and alpha and beta peaks from MEG using two large datasets (n=89 and n=189). We found that more robust anti-correlations between DMN and DAN regions are related to higher peak frequencies of alpha and beta rhythms in the brain. Subjects with higher alpha peak frequencies also show stronger positive within-network connectivity in both the DMN and DAN networks. Females show stronger correspondence between the two measures as compared to males. This association between two non-invasive cognitive neuroscience modalities adds to the growing literature on the biomarkers of cognitive and mental health.

**Key points:**

- MEG/EEG biomarker of brain health is correlated with rs-fMRI biomarker of brain health
- Alpha and beta peak frequency correlated with DMN-DAN functional connectivity
- Stronger within-network connectivity for subjects with a high alpha peak frequency
- Females have a stronger association between peak frequency and DMN-DAN connectivity

## 1 Introduction

Mental health disorders have become a pandemic with climbing rates of anxiety, depression and suicidal tendencies. There is an increasing need for biomarkers of mental and cognitive health. Recent advances in the field of cognitive neuroscience in general and neuroimaging, in particular, allow us to use non-invasive measures like Electroencephalography (EEG), Magnetoencephalography (MEG), functional MRI (fMRI) and diffusion MRI (dMRI) to make estimates about cognitive health (Woo et al., 2017; Dubois & Adolphs, 2016; Yamashita et al., 2018; Lefebvre et al., 2018; Hong et al., 2020; Murty et al., 2020; Cesnaite et al., 2023).

The Default Mode Network (DMN) plays an integral part in the intrinsic processing related to autobiographical information, long-term memory processes and social cognition (Buckner et al., 2008; Buckner and DiNicola, 2019; Andrews-Hanna, 2012; Spreng et al., 2014; Spreng and Grady, 2010, Buckner and Margulies, 2019; Margulies et al., 2016) and is an important network in determining the cognitive health of an individual (Coutinho et al., 2016; Mohan et al., 2016). Studies have shown that reduced connectivity within the DMN and between other regions and the DMN is associated with various disorders (Bozhilova et al., 2018; Kim et al., 2017; Mak et al., 2017; McCormick and Telzer, 2018; Mohan et al., 2016; Mowinckel et al., 2017; Querne et al., 2017; Zöller et al., 2017). The Dorsal Attention Network (DAN) on the other hand is related to extrinsic processing and is activated in attention-demanding tasks (Fox et al., 2005). ADHD, and attentional deficits are linked to departure from the normal functioning of the DAN (McCarthy et al., 2013; Spreng et al., 2014). DMN and DAN have been found to be anticorrelated (Fox et al., 2005) wherein cognitively demanding tasks activate the DAN and suppress the DMN (Greicius and Menon, 2004). Studies have found DMN connectivity with other networks to be a biomarker of cognitive health (Falahpour et al., 2016; Kim et al., 2017; Mowinckel et al., 2017; Soares et al., 2017) and expert meditators have a higher DMN-DAN anticorrelation as compared to healthy controls (Devaney et al., 2021).

Alpha waves, as detected using invasive and non-invasive measures, are the most readily observed brain waves. Alpha waves generate and propagate from parietal cortex to the sensory regions down the sensory hierarchy and are suggested to have a role in sensory feedback (Halgren et al., 2019). Studies have show a relationship between alpha band power and cognitive decline (Frutos-Lucas et al., 2018; López-Sanz et al., 2016; Poza et al., 2007), working memory capacity (Tran et al., 2016), and age (Frutos-Lucas et al., 2018; Garcés et al., 2013). Beta waves correspond to top-down signals generating from prefrontal cortex (Miller et al., 2018). Research has also shown the role of beta waves in cognitive health (Newson and Thiagarajan, 2019) and working memory (Cheng et al., 2022)

In order to investigate these questions, we examined two large, multi-modal datasets for which resting-state fMRI and MEG data is available for individual subjects. We analyzed the resting state functional connectivity between DMN and DAN across subjects from Human Connectome Project (n=89) (Van Essen et al., 2012) for whom both magnetoencephalography (MEG) and resting state fMRI (rsfMRI) were available. We found that the magnitude of anticorrelation between DMN and DAN is negatively associated with the peak alpha and peak beta frequencies as extracted from the MEG dataset in healthy subjects. High and low peak alpha frequency (PAF) subjects demonstrated stronger within network connectivity in the Dorsal Attention and Default Mode regions and stronger in-between Dorsal Attention and Default Mode Network connectivity. We also found that females have a stronger relationship between the measures as compared to males. We replicated our findings on another multi-modal dataset called MOUS (Mother Of all Unification Studies; n=189) (Schoffelen et al., 2019) with resting state data acquired during both modalities. Ours is one of the first studies to demonstrate the connection between the multimodal (MEG and rsfMRI) measures of cognitive health (Engemann et al., 2020; Vaghari et al., 2022; Xifra-Porxas et al., 2021)

## 2 Methods

Within-subject MEG and resting-state fMRI data were examined from two publicly available data sets, the Human Connectome Project (HCP) (Van Essen et al., 2012) and the Mother of All Unification Studies (MOUS) (Schoffelen et al., 2019).

### 2.1 Datasets

#### Human Connectome Project

The Young-Adult HCP acquired both MEG and fMRI datasets on each of 93 subjects. Out of the 93 subjects, 89 subjects (48 males) had completed full protocols in both paradigms and were selected for the present analysis. Informed consent was obtained from all subjects and the study was approved by the Washington University institutional review board. The dataset is publicly available on ConnectomeDB, the data management platform of the HCP (https://db.humanconnectome.org).

Each subject participated in two days of scanning in a custom Siemens CONNECTOM Skyra MRI Scanner. Along with high resolution structural T1 and T2 weighted MRI scans, four resting state fMRI runs of fifteen minutes each (60 min total), with eyes open were collected (Voxel resolution = 2 mm isotropic, 72 slices, multi band factor 8, TR = 720 ms, 1200 TRs) resulting in a total number of TRs of 4800 concatenated across the four runs. Subjects were asked to look at the visual fixation cross on a projected computer screen and do nothing in particular. The resting state data were collected across the two fMRI scan days and resting-state runs were interspersed with other task and structural acquisition runs in the HCP protocol. On a separate day, the subjects went through MEG acquisition with three runs of six minutes of resting state data acquired in a supine position with eyes open directed to a fixation cross. The MEG was collected in a shielded room on a whole head MAGNES 3600 (4D Neuroimaging, San Diego, CA) with 248 magnetometers, 27 reference channels and 64 EEG channels with a sampling rate of 2034 Hz. Head position was tracked using five localizer coils along with structural MR data and head surface tracings.

#### Mother of Unification Studies

MOUS acquired both MEG and fMRI datasets on each of 204 subjects. Some parts of the paradigms were missing for fifteen subjects, so 189 subjects (92 males) were included in the analyses. The dataset included structural, diffusion and functional data with resting state and task runs. These data were collected at Donders Centre for Cognitive Neuroimaging and are publicly available on the Donders Institute Repository (https://data.donders.ru.nl). The Ethics Committee at Radboud University approved the study. All subjects were informed and ethical informed consent was obtained.

MEG and fMRI data were collected on the same day, the order was counterbalanced across subjects. The resting state fMRI scan was acquired on a Siemens Trio 3 T Scanner (Voxel resolution 3.5 × 3.5 × 3 mm, TR = 1680 ms, 29 oblique slices, total 266 TRs) along with high-resolution structural scan and diffusion weighted runs. Subjects were asked to keep their eyes closed and not to think of anything in particular and not fall asleep during the seven minute resting-state scan. The MEG resting-state measurements consisted of 5 minutes of (eyes-open) data, collected using a 275-channel axial gradiometer CTF MEG system (CTF MEG International Systems, Canada) with a sampling frequency of 1200 Hz with an anti-aliasing low pass filter applied at a 300 Hz cut-off. Localizer coils attached at the nasion, left and right ear canals helped keep track of the position with respect to the MEG sensors. Head position was monitored and movements were generally within 5 mm of the original position. Further details of acquisition can be found in the dataset paper (Schoffelen et al., 2019). During the MEG resting state scan of five minutes, subjects were asked to fixate on a cross with eyes open and not to think of anything specific.

### 2.2 fMRI data analysis

The HCP data were preprocessed using the HCP minimal preprocessing pipeline (Glasser et al., 2013) and registered to surface space in the CIFTI grayordinates 32k format. The preprocessing included artifact correction, gradient non linearity correction, motion correction and EPI distortion correction,temporal denoising and high pass filtering at 100 s. As the acquisition was done at a fast TR of 720 ms, no slice time correction was done. The structural, T1w and T2w and functional images were registered from the subjects’ native structural space to MNI space and to the native surface mesh using the Freesurfer pipeline followed up by registration to high resolution 168k and finally to low resolution 32k CIFTI format.

For the MOUS dataset, we used the default fMRIprep toolbox pipeline (Esteban et al., 2019) for the preprocessing and registration of the subjects to fsaverage6 surface space. For the anatomical data preprocessing the T1-weighted image was corrected for intensity non-uniformity, skull-stripped, segmented into cerebrospinal fluid, white-matter and gray-matter and brain surfaces reconstructed using recon-all. For the functional preprocessing, each resting state run was co-registered (bbregister), slice time corrected, and resampled to fsaverage6 surface space. High pass filtering was done at 128 s. CompCor noise correction including head motion and other physiological regressors was applied, retaining components that accounted for 50 percent variability and dropping others. No spatial smoothing was done across the two datasets.

Functional connectivity analysis with both datasets employed ROIs defined by the Schaefer implementation (Schaefer et al., 2018) of the Yeo-7 network atlas (Yeo et al., 2011) with 100 parcels. We extracted the preprocessed time course signals for all the vertices in DMN and DAN ROIs and computed the mean resting state time courses across all vertices within each ROI. The ROI time courses were then correlated for all pairs of ROIs. We averaged the within hemispheric DMN-DAN ROI connectivity to give us a final measure of anticorrelation between the networks for a single subject. Correlations were computed using Pearson method as implemented in stats module of the SciPy toolbox in python (Virtanen et al., 2020)

We also computed rs-FC maps, using ROIs as seeds. We used the Schaefer 100 Parcel atlas to compute mean resting state signals for each ROI and computed the seed x seed map using Pearson correlation. The seed to seed maps were computed separately for all the subjects and also averaged separately for high and low alpha peak frequency subjects. The sign corrected difference map was computed between the high and low alpha subjects’ seed to seed connectivity map. During subtraction the sign of the correlation was preserved to demonstrate the difference in amplitude of correlations and anticorrelations in the sign corrected difference map.

A seed to vertex connectivity map was created for mPFC ROI. The map was created separately for all the subjects and averaged separately for high alpha (> 11 Hz) and low alpha (< 9.6 Hz) subjects. The sign corrected connectivity map was created in the same way as mentioned above, where the sign of the connectivity was preserved and amplitude difference calculated to highlight the difference in correlation and anticorrelation of all the vertices with the mPFC seed.

To find differences between the within and between network connectivity for high and low alpha subjects, we first averaged the time course signals within each ROI of the DMN and DAN networks, as given in the Schaefer-Yeo atlas. Each ROI averaged time course signal was correlated with each other. We averaged the within network connectivity using the correlation values in the ROIs separately for DMN and DAN and between network connectivity with ROIs from both DMN and DAN. We computed these values separately for both the hemispheres. We performed independent t-tests to compare the within and between network connectivity for high and low alpha subjects using SciPy toolbox (Virtanen et al., 2020) independently for the left and right hemisphere.

#### 2.3 MEG frequency analysis

We processed and analyzed MEG data in accordance with the recommendations of Gross et. al. (2014). The released MEG datasets were already visually inspected and manually checked and any epochs with motion artifacts were removed. Line noise was removed using a notch filter. 1 bad channel was removed from the MOUS dataset and 2 from the HCP. We did not perform the Independent Component Analysis (ICA) based signal removal as it also removed the alpha and beta wave components. We then removed the global signal from the data and then performed a band pass filtering between 2 and 45 Hz before computing the spectrogram.

In order to extract the peak frequency, amplitude at peak frequency, and band power, first we computed the frequency spectrogram of all the clean data from MEG channels and used the multi taper method as implemented in the multitaper_spectrogram package (Prerau et al., 2016). We set the time bandwidth as 4 seconds, number of tapers as 7 and a window size of 10 seconds with 2 second moving time step to compute the multitaper spectrogram. We averaged the spectrogram across the entire duration of the run to give us one spectrogram for the resting state data. We used the FOOOF toolbox (Donoghue et al., 2020) to extract the peak amplitude, frequency and band power in the alpha (8-13 Hz) and beta (13-30 Hz) frequency ranges. The number of peaks to detect was limited to three. Some channels that did not have alpha or beta band power, those values were set to null and not analyzed further.

To extract the relevant channel across the subjects, we averaged the peak amplitude and frequency and plotted it separately for the alpha and beta range on the brain map. We used mne python package for plotting the topomap (Gramfort et al., 2013). We chose bilateral parietal channels for alpha band and frontal channel for beta band based on the channels with high average peak frequency across subjects in the respective bands. Additionally, the channels were chosen based on prior studies that examined these regions thoroughly ((Pfurtscheller et al., 1997; Lundqvist et al., 2018; Fischer et al., 2018; Halgren et al., 2019). The peak frequency, amplitude and band power for these selected channels were then used in further analysis.

We also examined group level differences in rsFC between high alpha peak frequency subjects and low alpha peak frequency subjects. High alpha subjects (n=20) were defined as those having peak frequency in the selected channel greater than 11 Hz and low alpha subjects (n=20) were defined as those having peak frequency less than 9.6 Hz.

## 3 Results

For each individual participant, we extracted independent measures from both MEG and fMRI resting-state data. Using MEG data we computed (as shown in Figure 1A) the peak alpha-band frequency (PAF) and amplitude, after preprocessing, spectral analysis and 1/*f* calculation (Donoghue et al., 2020). Using resting-state fMRI data, we computed functional connectivity (as shown in Figure 1B) between the Dorsal Attention and Default Mode Network by calculating the mean resting state signal within the ROIs defined in the Yeo 7-Network atlas after preprocessing and registering to the surface space and taking a pairwise correlation between the signals. We utilized this metric as earlier studies have used the DMN-DAN connectivity as a measure of health (Lois & Wessa, 2016; Devaney et al., 2021) To identify the MEG channels of interest, we computed the peak amplitude and frequency for alpha and beta bands for all the channels and plotted these data on a topomap (Figure 2). Alpha band shows high amplitude and frequency in the central and parietal regions whereas the beta band shows high frequency in the central and frontal regions and high amplitude in the central and parietal regions. It suggests that the activity in the beta band decays from parietal to frontal but becomes faster in the frontal areas.

**Figure 1:**
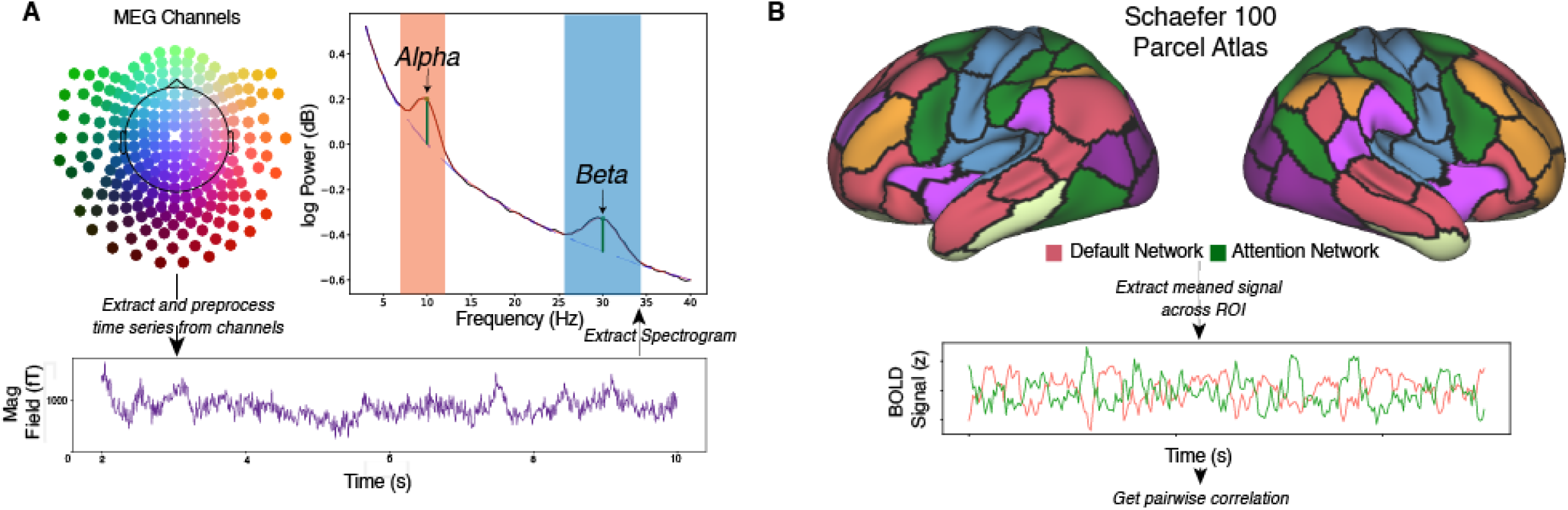
Method Description: We compute cognitive metrics from MEG and fMRI sources while subject undergo resting state. A) From MEG data, the raw time series acquired during resting state is extracted and preprocessed from relevant channels. We then extract the frequency spectrogram from the preprocessed time series. For alpha and beta bands, we extract the peak frequency and the amplitude of the frequency using FOOOF method (Donoghue et al., 2020). B) The same subjects undergo resting state in the MRI scanner, BOLD signals are extracted after preprocessing and registration to the surface space. We then compute the mean signal across ROIs derived from the Schaefer 100 atlas (Schaefer et al., 2018) which are then pairwise correlated to give us the resting state functional connectivity between the ROIs. Nodes of the Dorsal Attention Network (green parcels) and the Default Mode Network (red parcels) were a key focus of this analysis.

**Figure 2:**
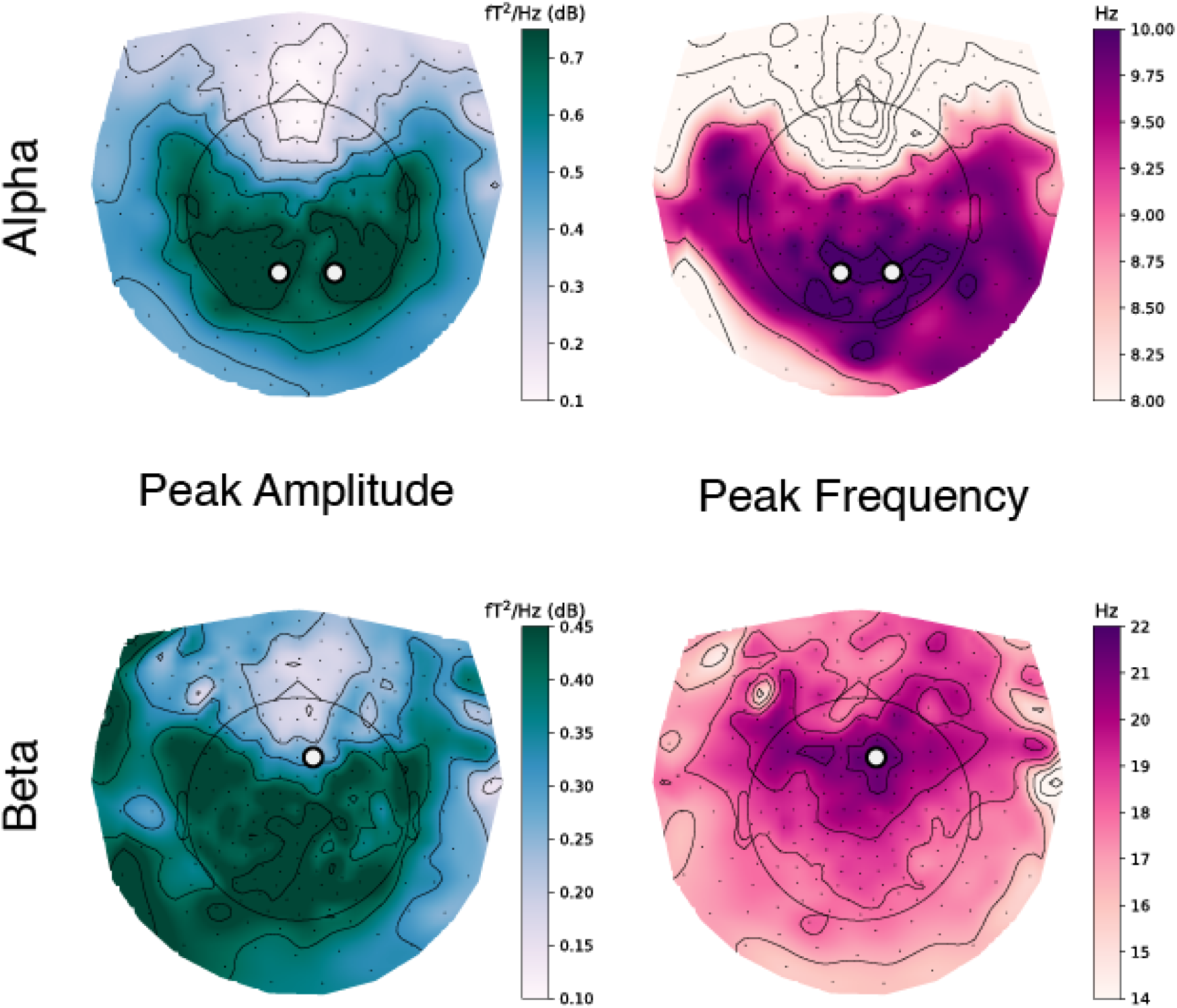
Peak amplitude and frequency topomaps: We compute the peak amplitude and frequency for the alpha and beta bands from all the channels averaged across the subjects from the MEG dataset as available in the HCP database and plot the values as topomaps. We then select the channels with high peak frequency (bilateral parietal for alpha and frontal for beta) for further analysis.

We narrowed our analyses to bilateral parietal channels (A103 and A107) for alpha rhythm and frontal A49 channel for beta rhythm. The channels were selected with high mean peak amplitude and/or frequency and were examined closely in prior invasive and non-invasive studies (Pfurtscheller et al., 1997; Lundqvist et al., 2018; Fischer et al., 2018; Halgren et al., 2019). We examined the relationship between DMN-DAN connectivity and peak alpha/beta frequencies (Figure 3) and observed a statistically significant relationship for both alpha (*r* =− 31, 95% CI [− 49, − 10], *t*(83) =− 2. 95, *p* =. 004) and beta channels (*r* =− 23, 95% CI [− 42, − 02], *t*(87) =− 2. 20, *p* =. 030). We explore the other relationships between alpha/beta band power, frequency and amplitudes in the Supplementary figures S1 and S2 and supplementary results section.

**Figure 3:**
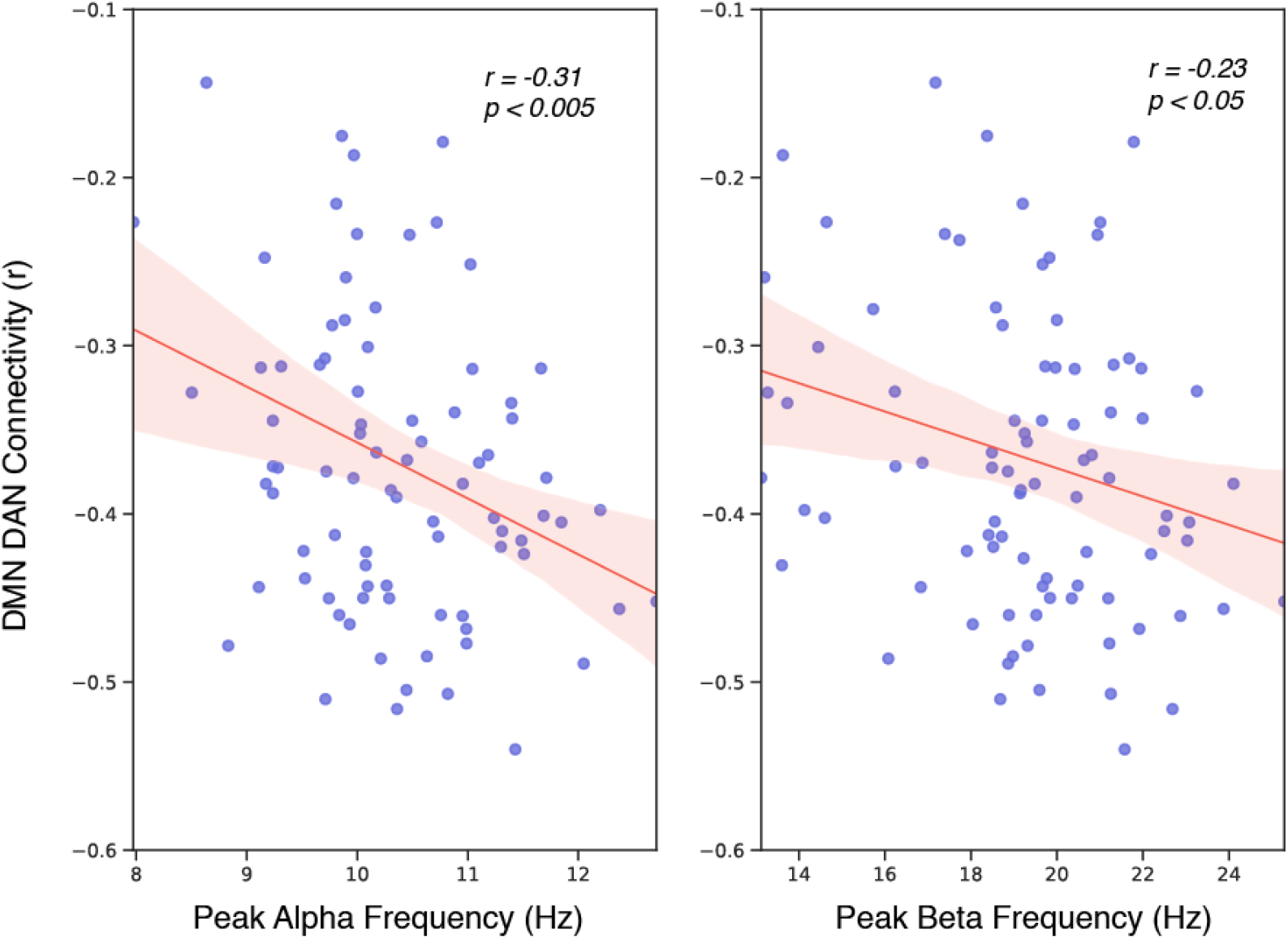
Relationship between Peak Frequency and DMN-DAN connectivity: In the HCP dataset, we find an anticorrelation between the subject specific peak alpha and peak beta frequency and the resting state functional connectivity amplitude between the Default Mode Network and Dorsal Attention Network. DMN and DAN are ‘opponent’ networks, so the strength of their connectivity is reflected in the magnitude of their negative connectivity (r). Therefore, higher peak alpha and beta frequencies positively correlate with the strength of the DMN-DAN opponency.

We separated our subjects into those with high frequency peak alpha (>11 Hz) and low frequency peak alpha (<9.5Hz) and analysed the seed to seed based connectivity matrices for all the 100 seeds in the Schaefer-Yeo 7 Network atlas (Schaefer et al., 2018) as shown in Figure 4B separately for high and low alpha subjects and also created a sign corrected difference map. We find that the within-network average functional connectivity for DAN and for DMN is higher for high alpha subjects (DAN: Δ*M* = 0. 04, 95% CI [0. 01, 0. 08], *t*(33. 46) = 2. 70, *p* =. 011 cohen’s d=0.62; DMN: Δ*M* = 0. 03, 95% CI [0. 00, 0. 05], *t*(37. 64) = 2. 15, *p* =. 038 cohen’s d=0.493 as shown in Figure 5. We also find that the between-network average connectivity for DMN and DAN is statistically significantly more anticorrelated (Δ*M* =− 0. 04, 95% CI [− 0. 06, − 0. 01], *t*(30. 20) =− 2. 59, *p* =. 015 cohen’s d=-0.595 for high alpha subjects. Looking closely at the networks, we find dorsal medial prefrontal cortex (dmPFC), posterior cingulate cortex (PCC) in the DMN and superior parietal lobule (SPL) contributes strongly to the connectivity. So, we analysed dmPFC closely as shown in Figure 4A. We computed the seed functional connectivity map which correlates the mean signal of the seed region and all the vertices on the cortical sheet. We plotted the seed correlation maps separately for the high and low alpha subjects in Figure 4A along with the sign corrected difference map between the two. The dmPFC map did not differ in connectivity within the DMN across the high and low subjects(Δ*M* = 0. 03, 95% CI [− 0. 01, 0. 06], *t*(35. 50) = 1. 47, *p* =. 149) but high alpha subjects had higher connectivity between the dmPFC and DAN (Δ*M* =− 0. 05, 95% CI [− 0. 09, − 0. 01], *t*(36. 68) =− 2. 82, *p* =. 008).

**Figure 4:**
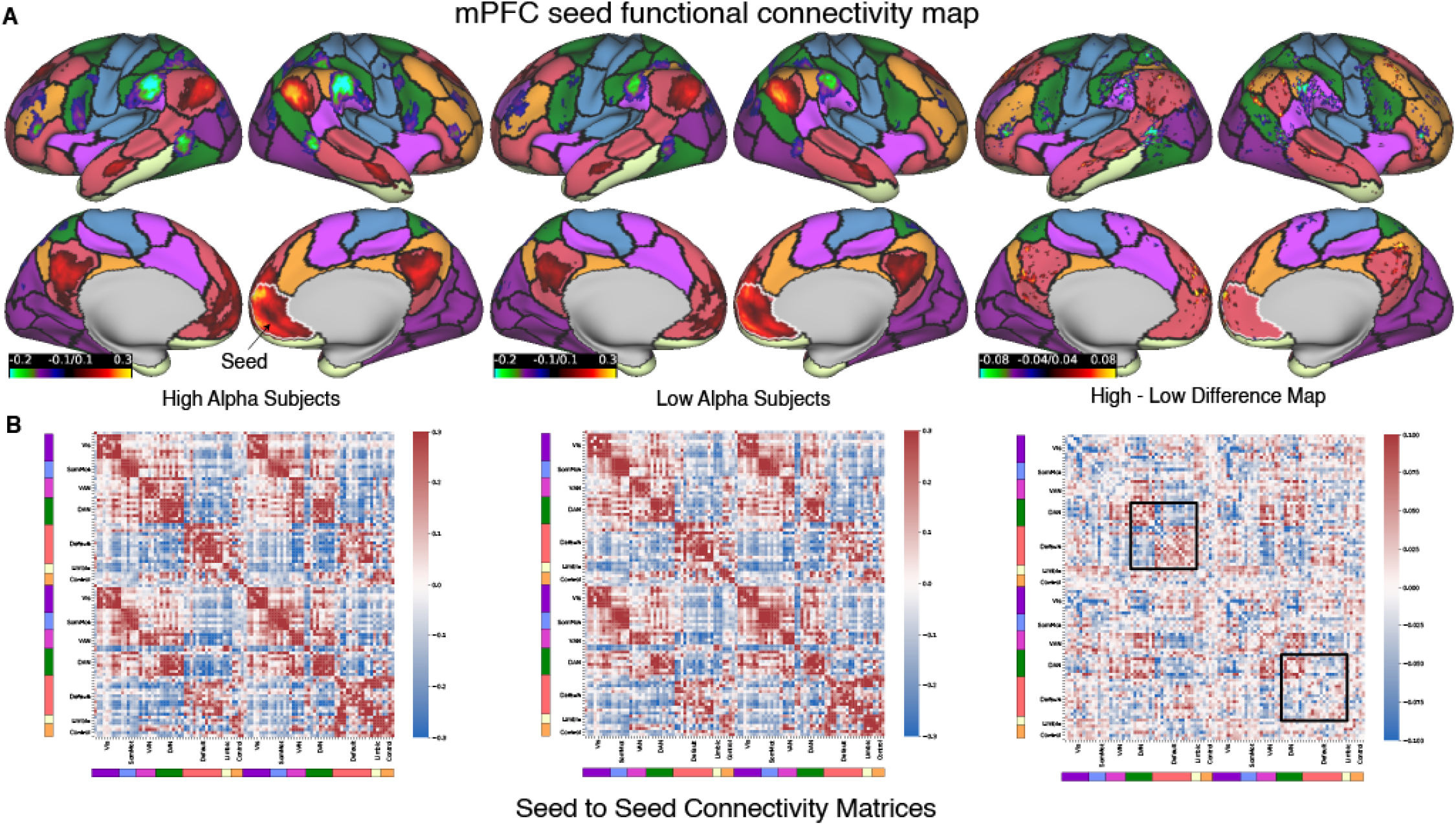
Functional Connectivity maps separated by High Alpha Peak Frequency and Low Alpha Peak Frequency subjects: A) We plot the dorsal medial prefrontal cortex (dmPFC) seed (white border) functional connectivity vertex wise map separately for subjects with high alpha (>11 Hz) and low alpha (<9.5 Hz) and create the sign corrected difference map between the two pointing out to the region having increased connectivity in the high alpha subjects as compared to low alpha subjects. B) We compute the seed to seed connectivity matrix where each seed belongs to one of the seven network as given in the Yeo 7 Network atlas (Thomas Yeo et al., 2011) and plot it separately for the high and low alpha subjects and compute the sign corrected difference map between the two. Black Squares highlight the within-hemisphere connections for DMN and DAN nodes.

**Figure 5:**
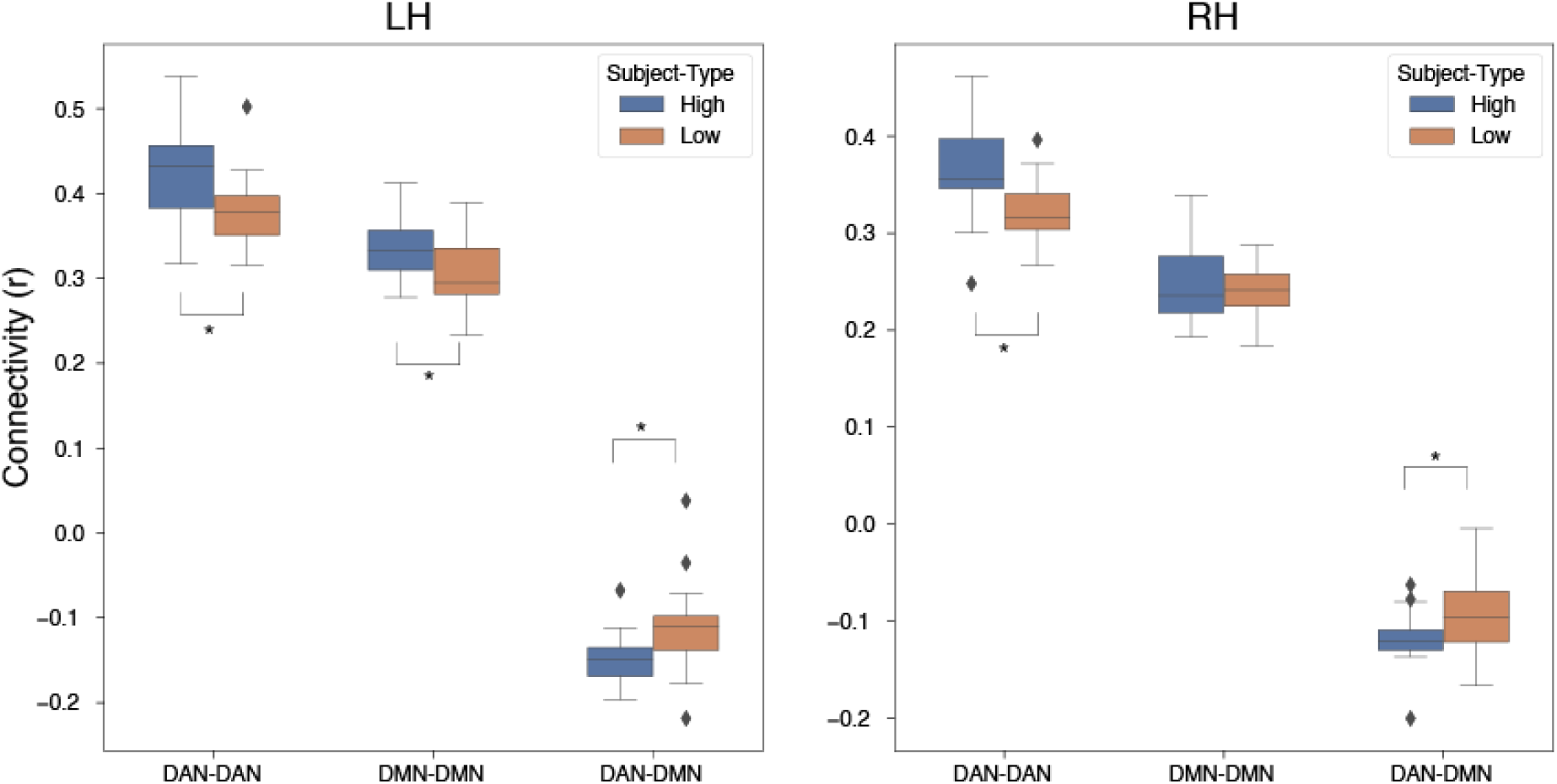
Within and between network analysis: High alpha subjects show stronger amplitude of within-network connectivity for Dorsal Attention and Default Mode networks and also between the two networks. * denotes statistical significance with p < 0.05.

We analysed the effects of sex on the PAF and DMN-DAN connectivity (Figure 6) and found out that females have a stronger anticorrelation between PAF and DMN-DAN connectivity (Pearson correlation: *r* =− 46, 95% CI [− 67, − 16], *t*(37) =− 3. 12, *p* =. 003) as compared to males (*r* =− 17, 95% CI [− 44,. 12], *t*(44) =− 1. 16, *p* =. 253). There were no difference in the peak alpha frequencies across the two groups (Δ*M* =− 0. 18, 95% CI [− 0. 59, 0. 22], *t*(77. 68) =− 0. 91, *p* =. 363). But the peak alpha amplitudes in the parietal channels is significantly higher for males (Δ*M* =− 0. 23, 95% CI [− 0. 38, − 0. 07], *t*(81. 29) =− 2. 87, *p* =. 005). We did not find any differences in the beta band activity between the two genders (Amplitude: Δ*M* =− 0. 02, 95% CI [− 0. 09, 0. 06], *t*(74. 39) =− 0. 47, *p* =. 643, Frequency: Δ*M* =− 0. 50, 95% CI [− 1. 63, 0. 62], *t*(86. 89) =− 0. 88, *p* =. 379).

**Figure 6:**
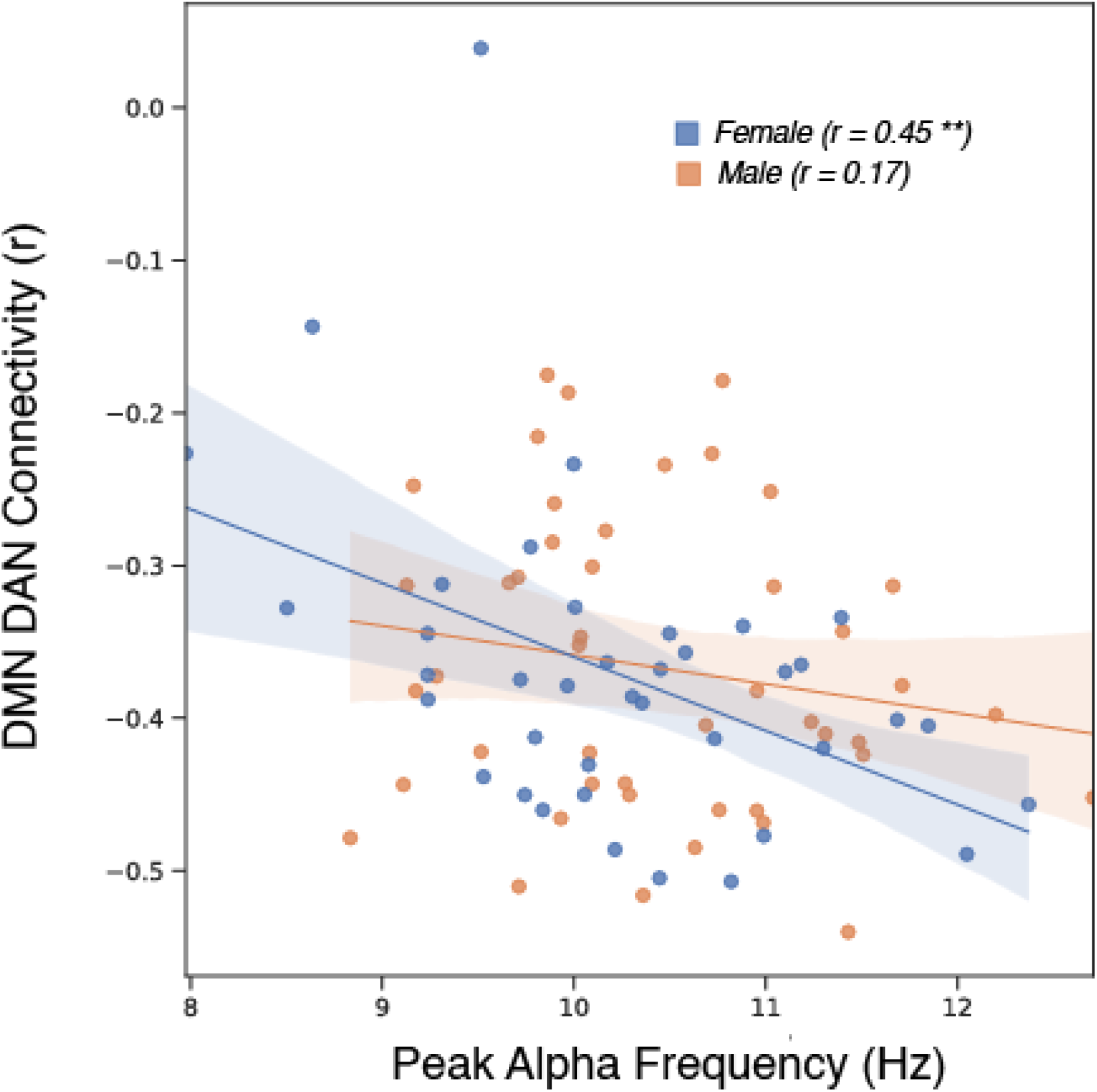
Gender differences in Alpha rhythms: We plot the relationship between the resting state functional connectivity between Default Mode and Dorsal Attention network and find that females have higher anticorrelation between the two measures as compared to males. * denotes statistical significance with p < 0.05, ** denotes statistical significance with p < 0.005.

We did not find any relationship between behavioral measures of memory and attention as computed using the NIH battery for the HCP subjects (all measures, p>0.05) for both DMN-DAN connectivity and the peak alpha/beta frequency and amplitudes. There were no differences in the PAF among the three age groups in the HCP dataset (21-25: M=10.41, S.D=0.76; 26-30: M=10.26, S.D=0.94; 31-35: M=10.39, S.D=0.94). The DMN DAN connectivity also did not show differences with age (21-25: M=-0.38, S.D=0.1; 26-30: M=-0.37, S.D=0.07; 31-35: M=-0.35, S.D=0.11).

In order to generalize and replicate our findings, we analysed the Mother of Unification studies dataset (Schoffelen et al., 2019) with 189 subjects. The resting state duration was 5 mins in the MEG and 7 minutes in fMRI with eyes closed, so the data was less in quantity as compared to HCP which affects the estimation of the true metric, we have seen in earlier studies that the connectome stabilizes after around 18 minutes of data (Tobyne et al., 2018) but still we were able to find similar pattern of activity for the peak amplitude and frequency [Figure 7] in the alpha and beta bands (stats?). We replicated the anticorrelation between the DMN DAN connectivity with the PAF (*r* =− 18, 95% CI [− 32, − 04], *t*(185) =− 2. 52, *p* =. 013) and peak beta frequency (*r* =− 19, 95% CI [− 32, − 05], *t*(190) =− 2. 61, *p* =. 010) and demonstrate that the two metrics are related.

**Figure 7:**
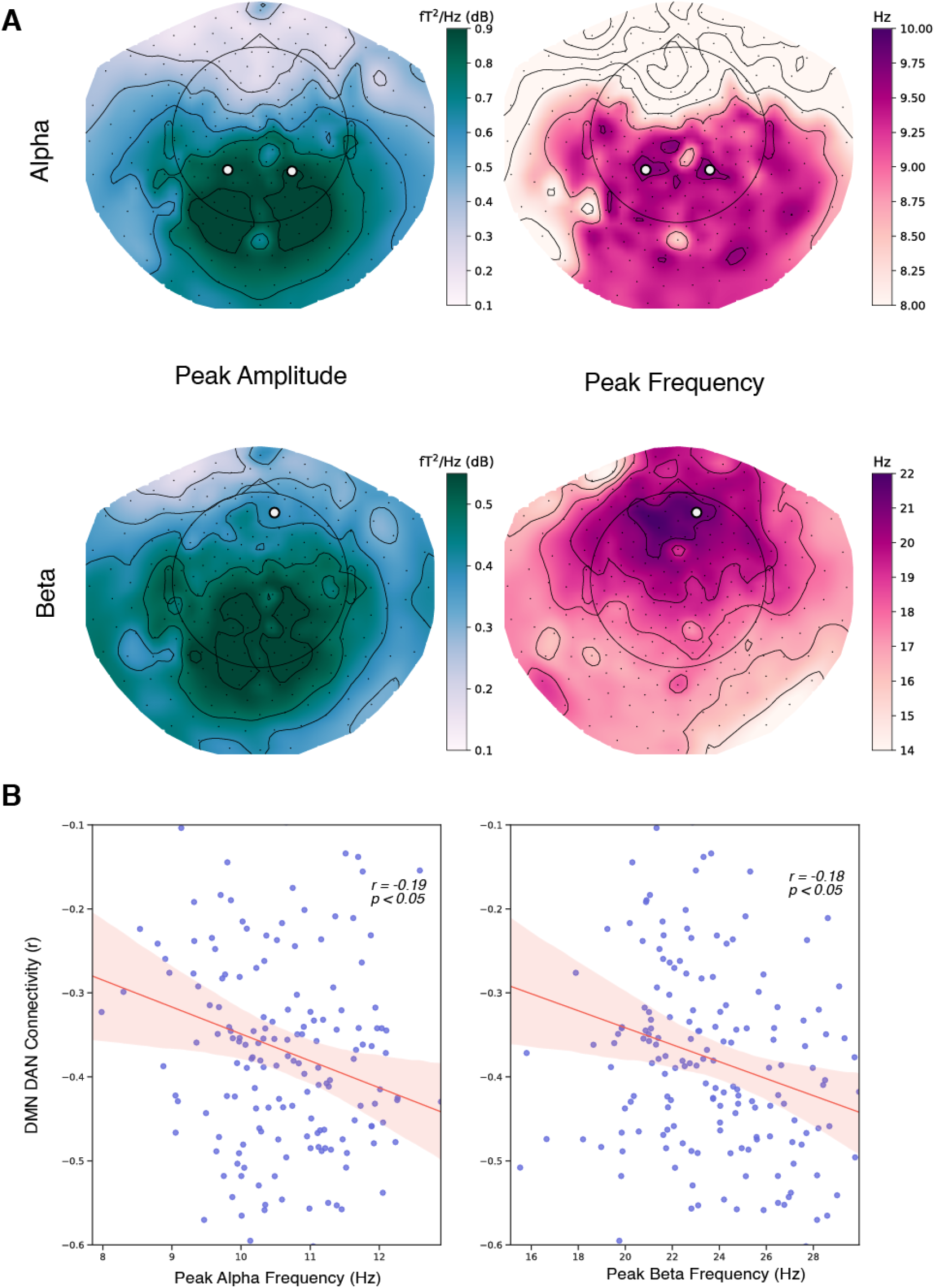
We replicate the analysis with another dataset called Mother of Unification Studies (MOUS, n=189) with both MEG and rsfMRI acquired for the subjects and found that we are able to replicate the findings with A) peak amplitude and frequency in the alpha and beta frequency bands and B) the relationship between DMN and DAN resting state functional connectivity and peak alpha and beta frequency. Although the quality and amount of data in the MOUS dataset was not better than the HCP dataset.

## 4 Discussion

We analysed the correlation between alpha and beta wave shift as detected in resting state MEG data and connectivity between DMN and DAN regions from resting-state fMRI data and found that the DMN DAN connectivity is negatively correlated with the peak frequency values in the alpha and beta range. We also found that subjects with higher alpha peak values had stronger within-network connectivity in the DMN and DAN regions. And we found gender differences between the alpha peak frequency and DMN-DAN connectivity, with females showing stronger correspondence between the two measures.

Default Mode Network was discovered as a region activated during baseline processing of the brain (Gusnard and Raichle, 2001; Raichle et al., 2001) and modulated by extrinsic attention and task demands (Greicius and Menon, 2004; Mckiernan et al., 2001). Later studies found that DMN and DAN make two anticorrelated networks (Fox et al., 2005; Golland et al., 2007). DAN is engaged during an attentive task and the degree of activation in the network relates to the task engagement. For some years, it was believed that the two are opposing networks and performance in a demanding task depended on how successfully a person can disengage the DMN. In the last decade or so, the role of the DMN has expanded from a task-negative network to one that is engaged in internal mentation (Buckner et al., 2008), intrinsic processing (Buckner and DiNicola, 2019; Jessica R. Andrews-Hanna, 2012) autobiographical memory (Spreng et al., 2014; Spreng and Grady, 2010), replay activation(Higgins et al., 2021), theory of mind (Saxe and Kanwisher, 2003), mind wandering (Christoff et al., 2016), and cognitive transitions (Crittenden et al., 2015; Smith et al., 2018; Tripathi and Garg, 2021).

DMN plays a key role in intrinsic tasks and disorders like AD, Attention Deficit Hyperactivity Disorder (ADHD), Parkinson’s, Epilepsy, and even stress and mood disorders can cause disruptions to DMN connectivity within and across networks (Bozhilova et al., 2018; Mohan et al., 2016; Mowinckel et al., 2017; Querne et al., 2017; Zöller et al., 2017). Depressive and anxiety states alter the DMN (Coutinho et al., 2016). Age influences DMN connectivity the strongest during the adult period and it weakens with age (Mak et al., 2017). For robust cognitive health, DMN connectivity within and across networks is essential (Kim et al., 2017; Mak et al., 2017; McCormick and Telzer, 2018).

Studies have found a relationship between the BOLD dynamics in DMN and alpha power (Gonçalves et al., 2006; Knyazev et al., 2011; Mo et al., 2013; Munck et al., 2008). Simultaneous EEG-fMRI studies show that regions like mPFC, PCC, and TPOJ in the DMN are strong for the alpha band (Marino et al., 2019) and alpha power is proportional to the activity in dorsal ACC, anterior insula and anterior PFC, all part of the DMN (Sadaghiani et al., 2012). A recent study on transcranial stimulation using alpha oscillations showed an increase of within network connectivity of the DMN (Clancy et. al. 2022). Studies that have created resting-state networks from MEG and EEG data consistently have found that the default mode network and the alpha band activity coincide (Brookes et al., 2011; De Pasquale et al., 2010; Jann et al., 2009). Alpha frequencies have been shown to mediate connectivity between DMN regions along with other bands like beta and theta (Samogin et al., 2019) and the alpha band DMN network has higher connectivity within and across the network (Samogin et al., 2020).

ECOG studies have shown that the alpha power is stronger in the somatomotor and default mode regions (Hacker et al., 2017). Alpha waves are generated in the parietal areas of the default mode (Hindriks et al., 2017) and dorsal attention networks and travel from higher association areas to early sensory areas (Halgren et al., 2019) suggesting hierarchical feedback processing or a top-down feedback loop. This line of research suggests an important role of alpha waves in cognition.

Beta band power is related to high DAN network connectivity (Samogin et al., 2020), cross-network connectivity between sensorimotor and core DMN networks (Sugata et al., 2020). Simultaneous studies show beta power from the posterior cingulate cortex and precuneus are positively correlated with BOLD response in the right frontal cortex (Neuner et al., 2014). Beta power is associated with cognitively demanding tasks (Fischer et al., 2018)

People with Subjective Cognitive Decline (SCD), Mild Cognitive Impairment (MCI), Dementia with Lewy Bodies (DLN) and Alzheimer’s Disease have decreased alpha power and PAF as compared to healthy age-matched adults (Garcés et al., 2013; López-Sanz et al., 2016; Peraza et al., 2018; Poza et al., 2007; Wiesman et al., 2021). Reduction in alpha power precedes change in PAF (López-Sanz et al., 2016). Subjects with a genetic disposition to AD (carriers of APOE4 gene) showed reduced PAF but physical activity improved the PAF pointing out the cognitive association of the measure (Frutos-Lucas et al., 2018). Concussion causes increased DMN and motor network connectivity across alpha through gamma frequency-based networks as determined using MEG (Dunkley et al., 2018). Sleep deprivation alters alpha power and causes inhibition of precuneus and PCC of DMN and the effect is not completely reversed with recovery sleep (Wu et al., 2021). Age has been shown to slow down PAF and alpha power (Tran et al., 2016). Another study showed although in a smaller group than females have higher PAF than males (Garcés et al., 2013). A study on meditators found the dynamic interplay between DMN and frontoparietal regions in the alpha band as the meditators transitioned from resting state to one of two different meditations (Marzetti et al., 2014). Meditators have higher anticorrelated DMN-DAN connectivity as compared to healthy controls (Devaney et al., 2021) suggesting meditation as an intervention for maintaining cognitive health.

Alpha power is associated with cognitive measures (Grandy et al., 2013). PAF is positively associated with general cognitive measures suggesting people with higher alpha power have improved cognition. Task demands also affect the PAF, a study found that individuals’ PAF was higher during demanding 2-back working memory tasks as compared to the 0-back conditions (Haegens et al., 2014). Children with Autism Spectrum Disorder (ASD) have lower PAF as compared to typically developing children (Dickinson et al., 2018). There is within-subject variability in the parietal and occipital PAF and larger across-subject variability in the measure (Quinn et al., 2021) suggesting that a broader window than 8-12 Hz is preferred for extracting alpha wave measures. A tACS-EEG study found a causal relationship between PAF and double-peak illusion (Cecere et al., 2015) where subjects report hearing the number of flashes paired with beeps. If beeps were closer than 100 ms, subjects reported two flashes when only one was presented. tACS stimulation on, above and below the PAF decreased and increased the temporal window. Higher PAF and smaller the temporal window, the lesser the perception of the flash suggesting alpha oscillations act as a “temporal unit of hierarchical visual processing.”

Alpha is generated in the default regions as evidenced by the multiple studies here, it acts as a top-down feedback wave (Cecere et al., 2015; Halgren et al., 2019) which is influenced by the state of alertness, awareness and disease. Patients with SCD, MCI and AD amongst others have shown a reduction in the alpha power (Garcés et al., 2013; López-Sanz et al., 2016; Peraza et al., 2018; Poza et al., 2007; Wiesman et al., 2021) and DMN connectivity (Mohan et al., 2016) suggesting slowing down of hierarchical processing due to disease. PAF is associated with cognitive health and is affected by age and cognitive decline. Interventions like meditation can prevent cognitive decline due to a reduction in feedback processing and stimulation interventions can be designed that can prevent the onset of the decline. This is one of the first studies to directly look at the correlation between two cognitive measures in a large cohort of subjects and replicate the finding across datasets. Though the current study is correlational, future stimulation studies coupled with concurrent EEG-fMRI could clearly establish a causal evidence between the alpha and DMN-DAN measures.

## 5 Conclusions

We investigated the role of resting-state functional connectivity between DMN and DAN regions and its association with alpha and beta peak values using MEG resting-state data. We found that alpha and beta peaks were correlated with the anti-correlated rsFC between DMN and DAN regions. Subjects with high alpha values are stronger within DMN and DAN network connectivity and in between network connectivity. Females have stronger relations between the two measures. The switching between DMN and DAN networks is important for cognitive health and alpha and beta peaks might be associated with such mechanisms. Ours is one of the first studies to show the association between the two suggesting that the network property is evident through multiple modalities and may result in the development of a more accurate index of cognitive wellness.

## 6 Acknowledgements

We would like to thank the WU-Minn Consortium (Principal Investigators: David Van Essen and Kamil Ugurbil; 1U54MH091657) for the Human Connectome Project database which is funded by the 16 NIH Institutes and Centers and by the McDonnell Center for Systems Neuroscience at Washington University. We’d also like to thank the Max Planck Institute for Psycholinguistics and Donders Institute for Brain, Cognition and Behaviour for the MOUS dataset.

## Data Availability Statement

Human Connectome Project (HCP) is publicly available on ConnectomeDB, the data management platform of the HCP (https://db.humanconnectome.org). The Mother of Unification Studies (MOUS) dataset is publicly available on the Donders Institute Repository (https://data.donders.ru.nl).

