## Supplementary figures for "Default Mode and Dorsal Attention Network functional connectivity associated with alpha and beta peak frequency in individuals"

### **Supplementary Results**

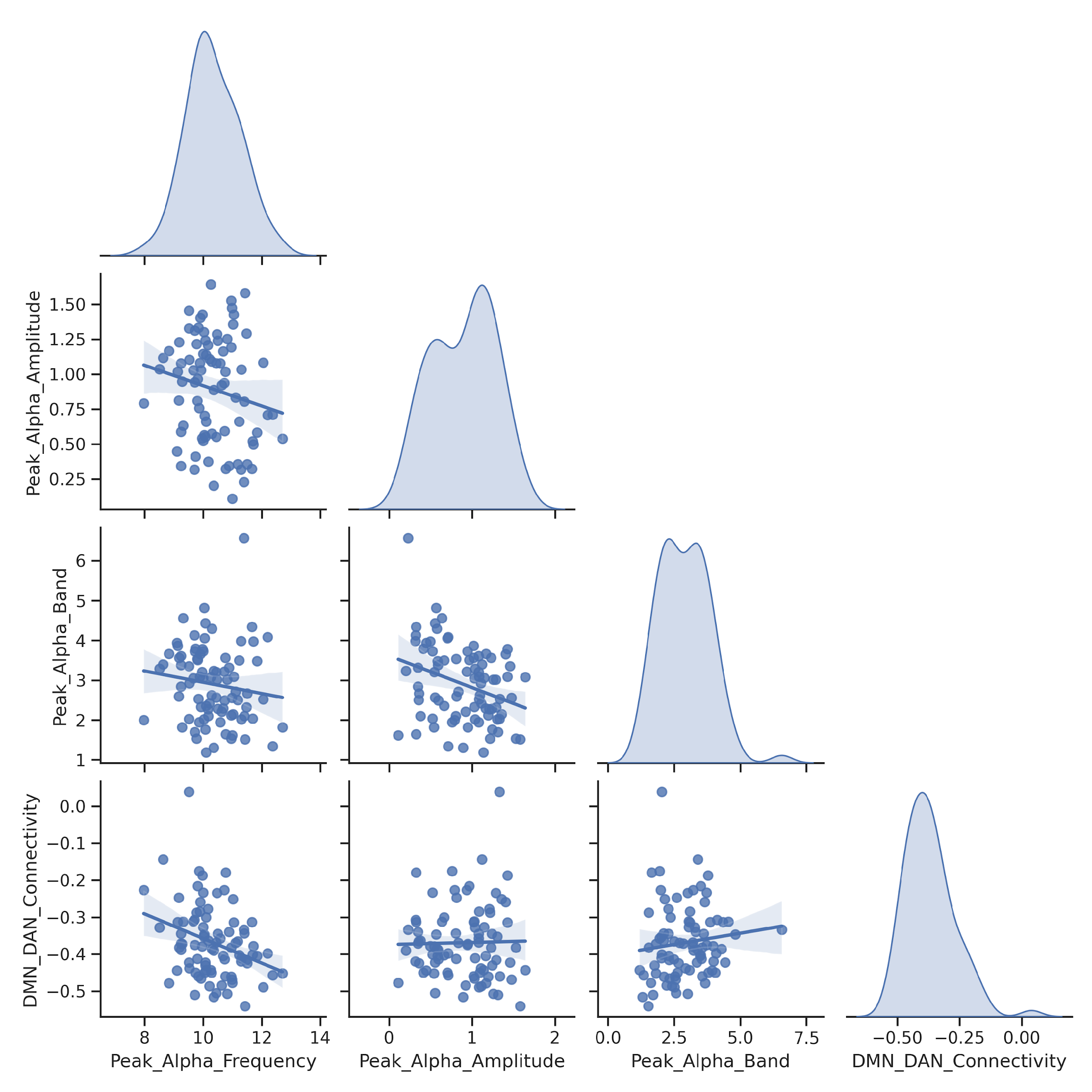

*Figure S1: Relationships among peak alpha frequency, amplitude, band power, and DMN-DAN connectivity for the bilateral parietal channels in the HCP dataset. The amplitude and the band power decreases as the peak alpha frequency increases.*

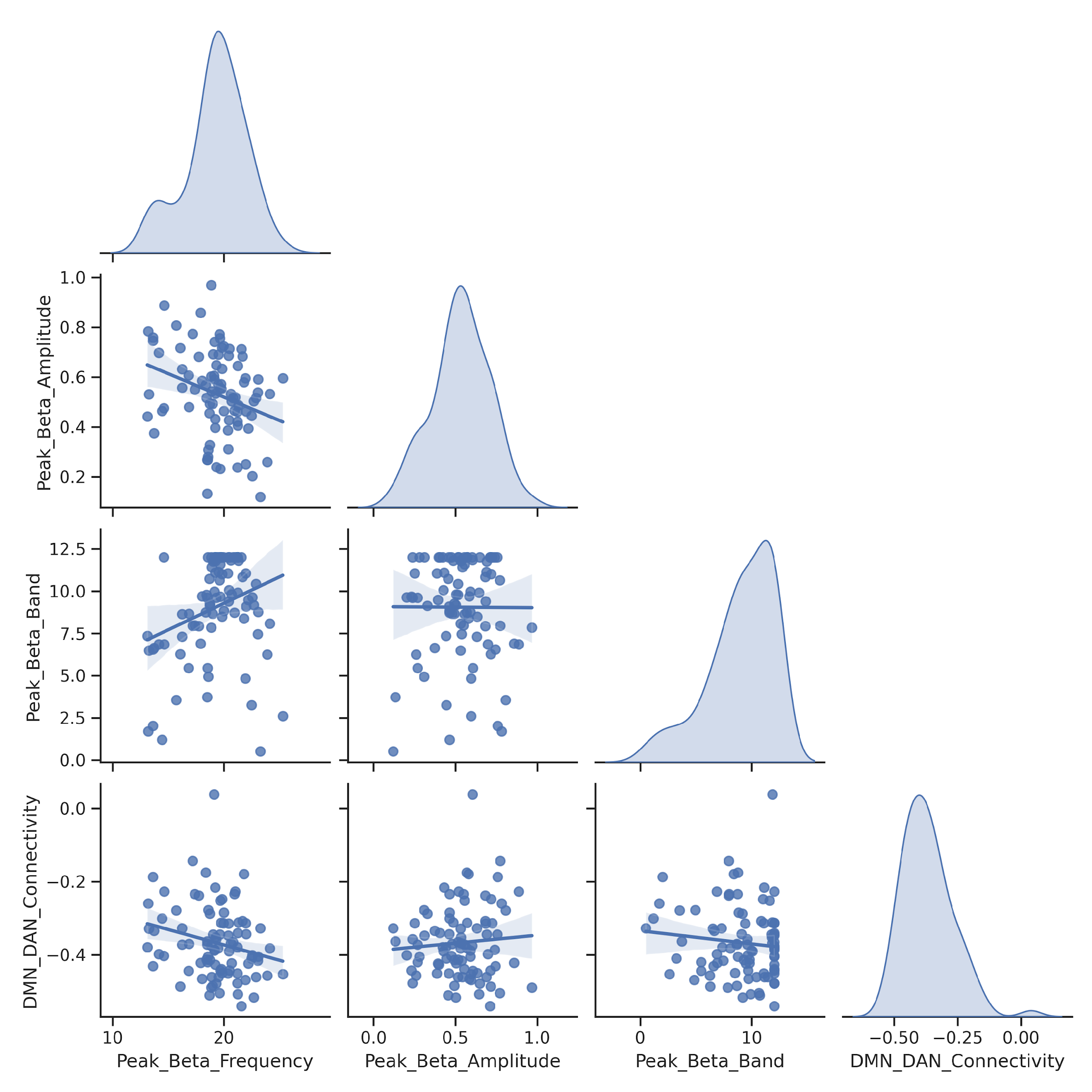

*Figure S2: Relationships among peak beta frequency, amplitude, band power, and DMN-DAN connectivity for the frontal channel in the HCP dataset.*

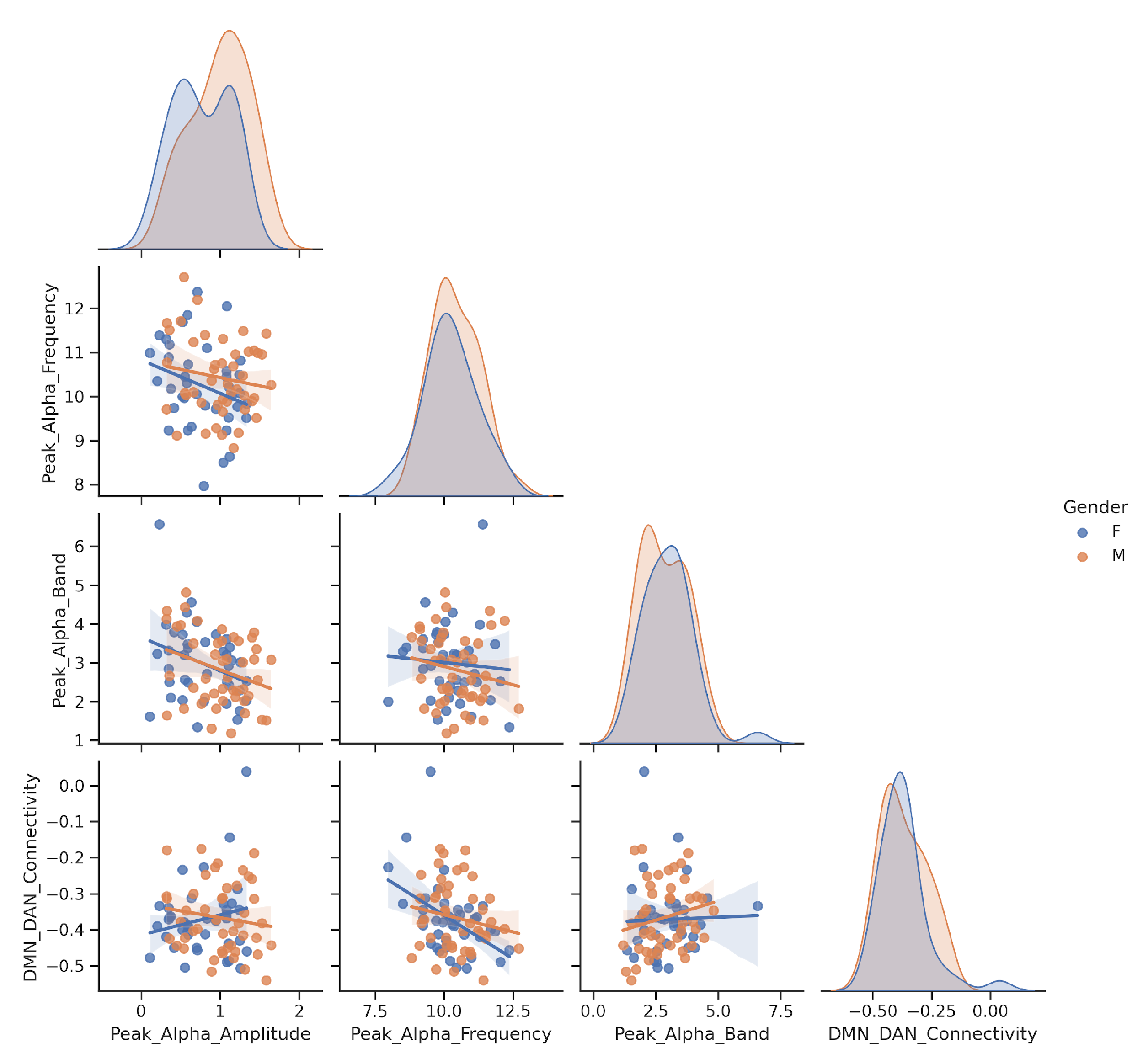

*Figure S3: Separating out the relationship among peak alpha frequency, amplitude, band and DMN-DAN connectivity for the frontal channel in the HCP dataset for the two genders.*

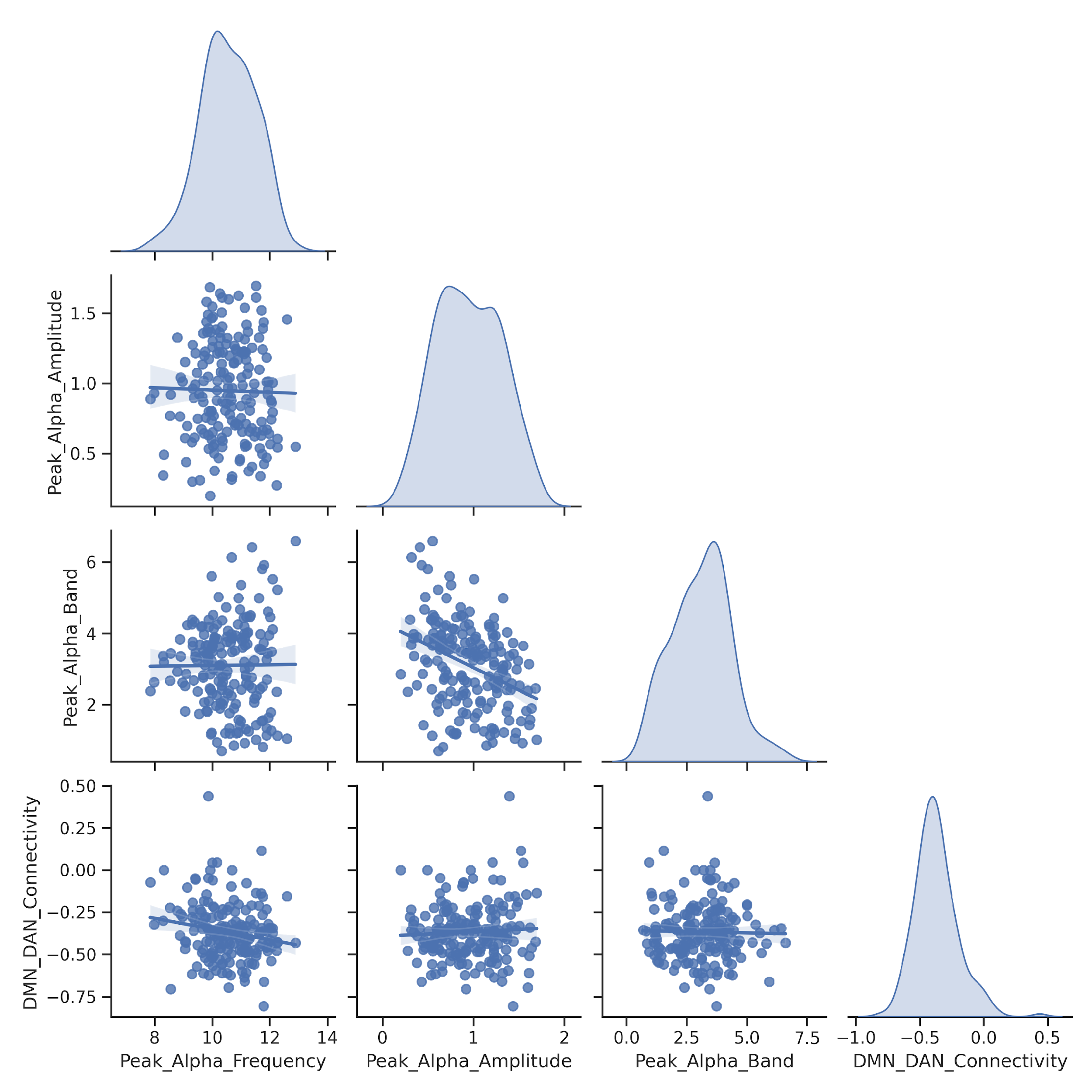

*Figure S4: Looking at the relationship among peak alpha frequency, amplitude, band and DMN-DAN connectivity for the bilateral parietal channels in the MOUS dataset.*

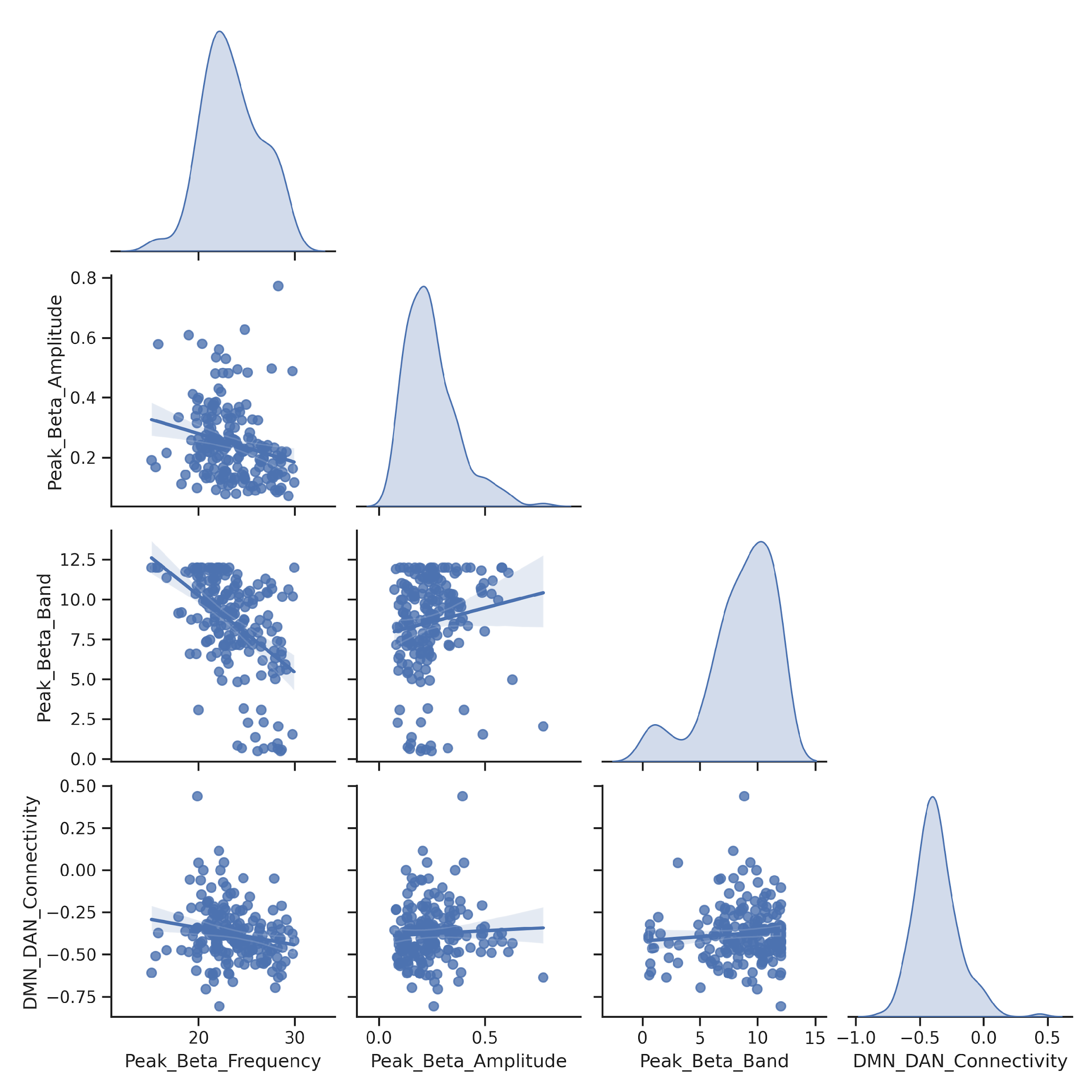

*Figure S5: Looking at the relationship among peak beta frequency, amplitude, band and DMN-DAN connectivity for the frontal channel5 in the MOUS dataset.*
